# Carbonate dissolution without acid: carbonate hydrolysis, catalyzed by photosynthetic microorganisms, in deteriorating stone monuments

**DOI:** 10.1101/2023.07.14.549033

**Authors:** Henry J. Sun, Gaosen Zhang, Zakaria Jibrin

**Author notes:** Key Laboratory of Extreme Environmental Microbial Resources and Engineering, Gansu Province; Northwest Institute of Eco-Environment and Resources, Chinese Academy of Sciences, Lanzhou, 730000, China (Gaosen Zhang).

## Abstract

Rapidly deteriorating stone (marble) monuments are generally blamed on acid rain. We suggest an alternative, not necessarily exclusive, culprit: rock-inhabiting cyanobacteria and microalgae, which may enter via fissures and dissolve carbonates as they propagate under the translucent stone surface. Cyanobacteria and algae absorb CO_2_ and HCO_3_ ^−^ and, in so doing, catalyze the reaction between carbonates and water itself. The resultant calcium hydroxide is expected to react with atmospheric CO_2_ and with adhered dust on its way out. We tested this hypothesis at the Forbidden City in Beijing, China, by inspecting stone monuments (dolomitic marble) for telltale signs of colonization and by studying an exfoliation crust with molecular and imaging techniques. The results, reported here, are consistent. Deterioration began in, and spread from, stone joints, cracks, and shattered stone edges. A cyanobacterial biofilm visible to the naked eye was present under the deteriorating stone surface. Colonized mineral grains were dissolved in a surface-controlled manner, i.e. along crystallographic and twinning planes. Secondary calcite, as well as clay minerals, were detected in the crust.

## Introduction

Rapidly deteriorating marble monuments have been documented around the world [1–12]. In the case of coarse-grained marbles, the cause is known. In marbles mineral grains, which are either calcitic (CaCO_3_) and dolomitic [CaMg(CO_3_)_2_], are interlocking. Such minerals are also anisotropic. When heated, they expand in different rates along different crystallographic axes [13–19]. The resulting strain to rock fabric is proportional to grain size: the larger the grains, the greater the strain [4, 12, 20, 21]. Repeated exposure to heat-cool cycles, therefore, results in stone fatigue and failure.

In the case of fine-grained marbles, the cause remains elusive, however. To be sure, observed changes in the optical property of deteriorating marbles implicates chemical dissolution. Pristine marble looks dull. Deteriorating marble looks transparent and reflective as if it is covered in sugar crystals, indicating that the mineral grains have been dissolved along their crystallographic planes [1, 6]. According to conventional wisdom, “sugaring” indicates exposure to acid or proton [1, 6, 22]. Thermodynamic considerations suggest otherwise, however. Detachment of ions from mineral surface is fast when by proton and carbonic acid, ceding the control of mineral dissolution rate to how fast dissolved ions are transported away from the mineral surface [23, 24]. Crystal corners, which stick out into undersaturated solvent, dissolve first, resulting in rounded crystals, the exact opposite of sugaring [23–26]. Correctly, sugaring should be interpreted as evidence of surface-controlled mineral dissolution. Surface control, in turn, requires the dissolution environment to be CO_2_ and proton deplete, so that the slow acting water, rather than carbonic acid and proton, is the dominant forward reaction [23, 24]:

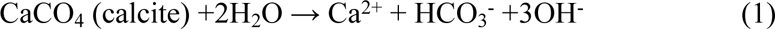

Carbonate hydrolysis is slow and reversible, however, so we need a catalyst. Two early studies described the behavior, though not the identity, of such a catalyst. While inspecting the Acropolis of Athens, Greece, Sofianopoulos [11] noted deterioration patterns that are difficult to explain unless one assumes that it is catalyzed by an airborne substance that has affinity for stone joints. More recently, Charola et al. [1] observed simultaneous casehardening and sugaring on a historic building in New York City. This places the site of mineral dissolution below, not on, the stone surface and indicates that the dissolved calcium is basic, not acidic. To summarize, whatever the catalyst is, it is ubiquitous in the environment; it is microscopic and readily airborne; it has an affinity for cracks and fissures; it propagates under the stone surface; and it absorbs the bicarbonate from carbonate hydrolysis but leaves the calcium hydroxide untouched.

The elusive catalyst, we argue, is endolithic cyanobacteria and microalgae. Their existence is well-documented in rocky deserts, of sandstones as well as marbles and limestones [27–34]. Porous rocks absorb rain, snowmelt, and dew [35–37]. With no canopy or epilithic lichens to intercept sunlight, sufficient photon fluxes are available near the translucent stone surfaces [38, 39]. HCO_3_ ^−^, as well as CO_2_, is a source of inorganic carbon, a metabolism that generates OH^−^ waste [40, 41]. The resulting CO_2_- and HCO_3_ ^−^-deplete water at the mineral-microbe interface is no ordinary water. In geochemical parlance, such water is said to be undersaturated with respect to carbonates and capable of accelerating carbonate hydrolysis [23–26]. Simply put, a photosynthetic microorganism cannot grow in a stone without stimulating its dissolution.

It is evident from equation (1) that a colonized limestone and dolostone would exude Ca(OH)_2_ and CaMg(OH)_4_, respectively. The exudate would be rinsed off if the stone surface is hydrated by snowmelt or wicked off if the stone is in contact with moist sediment. Absent these conditions, the exudate would react with atmospheric CO_2_ and precipitate calcium carbonate on the way out of the stone, producing casehardening or encrustation. The exudate, being alkaline, is also expected to dissolve adhered silicates dusts, supplying dissolved elements for clay mineral formations in the stone [42].

The resulting aggressive stone surface reduction, as well as the role of stone joints in initiating it, can now be explained also. The rapid surface reduction is due to mineral dissolution by se. Colonized stone ultimately loses cohesion, causing the overlying stone surface, about 1-2 mm thick, to exfoliate [28]. The remaining organisms grow deeper, while the exfoliated organisms are propagules, dispersed by wind over vast geographic distances, possibly globally [43, 44]. These propagules are deposited on all surfaces, but only cracks and fissures may provide a foothold and support chasmolithic growth [31, 45].

Below, we present a case study in the courtyards of the Forbidden City and Temple of Heaven in Beijing, China. Here, a fine-grained white dolomitic marble, in the form of a temple, totems, decorative pathway stones, stairs, and rails, is in an unshielded outdoor environment for more than 600 years [46] and, like ancient stone monuments elsewhere in the world in such setting, show signs of rapid deterioration. We examined deteriorating stones for characteristic texture, color, and association with cracks and fissures. We also analyzed a small exfoliation crust with molecular and imaging techniques. All results are consistent with the above-described microbially catalyzed process, none with acid rain.

## Material and Methods

Genomic DNA was extracted using the UltraClean® soil DNA isolation kit (MoBio Laboratories, Solana Beach, CA). Bacterial 16S rRNA genes were PCR-amplified using the primer set 27F and 1492R and annealing temperature of 50°C. PCR product was purified using the UltraClean® GelSpin DNA purification kit (MoBio Laboratories, Solana Beach, CA) and cloned into the vector pCR2.1 TOPO TA (Invitrogen). Unique clones were sequenced by Functional Biosciences (Madison, WI) using the primer set M13F-20 and M13R-27. Nucleotide sequences were screened for chimeras using Bellerophon and verified manually (Huber, Faulkner et al. 2004). Closely related sequences in the GenBank database were identified by BLAST search (Altschul, Gish et al. 1990). Similarity analyses were performed using FASTA. Sequences were aligned using multiple-alignment CLUSTAL W 2.0 software (Larkin, Blackshields et al. 2007). Phylogenetic dendrograms were generated using the neighbor-joining method (Saitou and Nei 1987). Tree topologies were evaluated by performing bootstraps analyses (Felsenstein 1985) of 1000 resamplings using MEGA 3.1 [47].

For scanning electron microscopy, samples were prepared in two ways. For studying stone-microbe relationship, the protocol of Dong et al. [48] was used. Briefly, specimens were rehydrated, fixed first in 0.5% formaldehyde and then in 1% glutaraldehyde, each for 15 minutes, and dehydrated in an ethanol series 15%, 30%, 50%, 75%, 95%, 100%, each for 15 minutes. After two additional changes in anhydrous ethanol, specimens underwent two soaks in hexamethyldisilazane (HMDS), each for 30 minutes. After decanting HMDS, specimens were air-dried and sputter-coated with gold. For EDS elemental analysis, marble embedded in epoxy resin was polished and sputter-coated with gold. Both samples were studied under a JEOL JSM-5610 scanning electron microscope equipped with an Oxford ISIS energy dispersive spectrometer.

Light microscopy examination of isolated cyanobacteria was done using a Zeiss Axioskop 2 epifluorescence light microscope. Petrographic thin sections were prepared by Burnham Petrographics LLC (Rathdrum, Idaho).

## Results

### Weathering patterns

The presence of an endolithic colony could be recognized by an altered stone surface and, in cases where the colony is young, by its association with cracks and fissures as well [28, 49]. In Figure 1 we include a photograph of each diagnostic feature in deserts, against which weathered stone monuments could be evaluated without destructive sampling. Figure 1A is a carbonate-cemented sandstone in the Mojave Desert that is partially colonized [49]. The uncolonized parts are black, due to the presence of a thin coat of ferromanganese oxides. This rock coat, known as desert varnish, formed before this boulder split from a vertical cliff and fell to the ground. In deserts, a smooth cliff is extreme arid because it may not be covered in snow, and rain only leaves a thin film of liquid water, which evaporates quickly. Under such condition a small amount of iron and manganese is extracted from adhered aeolian dust and precipitated on the rock surface [50, 51]. In its current tilted position, the stone’s moisture condition is improved, but only to the point to support endolithic lichens, not epilithic lichens. Colonization began in cracks and fissures and subsequently spreads under the varnished stone surface, causing it to exfoliate bit by bit. The resulting white, receded patches reflect where the cracks and fissures used to be.

**Figure 1.**
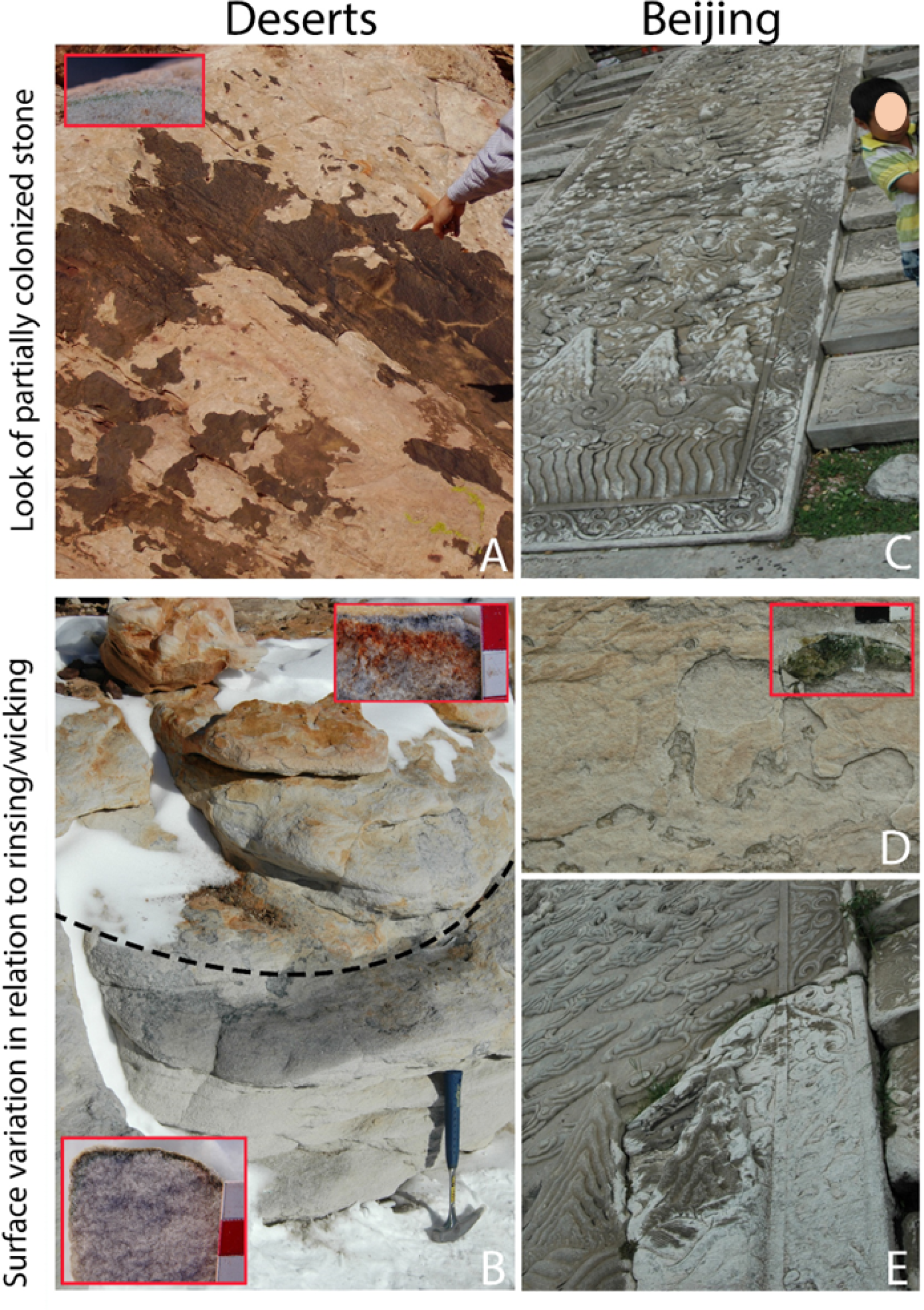
Signs of a colonized stone. Crack/fissure-centered weathering patterns on a sandstone in the Mojave Desert (A) and on a pathway stone in a courtyard at the Forbidden City (B). Exfoliation crusts on the high and dry upper part of a sandstone slope in Antarctica (above dashed line) and on a vertical wall in Beijing, versus grain-by-grain disintegration or “sanding” on the relatively moist lower half of the sandstone in Antarctica (below dashed line) and on a pathway stone in Beijing.

Figure 1B is a silica-cemented sandstone in the high-altitude areas of the Antarctic dry valleys region, included to illustrate how the hydrological environment controls stone texture and color. In this high-land polar desert, snow is the only form of precipitation, and the prevailing wind, which blows from the south, in the photograph towards the viewer, frequently sweep the highland [36, 37]. As a result, most of the snow on the upper half of the slope is blown off before it has a chance melt in the summer, when insolation causes rock temperatures to rise above 0°C [36]. Consequently, this part is relatively arid and colonized by endolithic lichens. The dissolved silica and iron from the colonized zone are reprecipitated on the stone surface as well as in the stone interior, resulting in an exfoliation crust and an accumulation zone, respectively (Figure 1B, upper inset). Snow on the lower half of the stone is, contrast, somewhat shielded from wind, this surface being leeward. It also absorbs snowmelt from sediment on the ground. The higher moisture content favors endolithic cyanobacteria, and the dissolved silica and iron are rinsed and wicked off, resulting in a bleached and crustless stone surface is. This surface recedes continually by dislodgement of individual sand grains, as opposed to a 1-2 mm thick crust in the upper part of the stone (Figure 1B, lower inset).

The same characteristic features were observed on weathered marbles in the courtyards of the Forbidden City and Temple of Heaven. Like the partially colonized sandstone in the Mojave desert, weathered surfaces were centered around cracks, shattered stone corners and edges, or bas relief (Figure 1C). Like the sandstone in Antarctica, altered marble texture and color is controlled by whether or not the stone has the possibility of being covered in snow. Beijing is in a cold temperate climate where about half of the precipitation is in snow. Although both snowmelt and rain activate the organisms, only a slowing melting snow cover rinses the stone surface while the organisms are dissolving it. Snow may collect on pathway stones and stairs, but not on vertical walls. Hence, like the upper part of the sandstone slope in Antarctica, weathered vertical walls were covered in exfoliation crusts that incorporate iron and redden over time (Figure 1D). Weathering damages on pathway stones, in contrast, resembled the lower half of the Antarctic sandstone. They had the texture of sandpaper and were snow-white as if they were bleached (Figure 1E).

Cyanobacteria were present underneath exfoliation crusts (Figure 1A, inset). Black-pigmented cyanobacteria were present in between mineral grains in sugared marble surfaces, consistent with the fact that cyanobacteria exposed to ultraviolet rays (UV) contain scytonemin, a brownish UV screening pigment in their polysaccharide sheaths [52, 53].

### Analyses of exfoliation crusts

A fractured exfoliation crust was imaged under the scanning electron microscope (SEM). The lower half of the crust was infiltrated by a biofilm, which grew in between and around mineral grains (Figure 2). The uncolonized outer layer shows what the colonized zone used to be: consisting of interlocking, smooth mineral grains. A petrographic thin section of the exfoliation crust was examined under a polarized light microscope to visualize the dissolved mineral grains in silhouette. The mineral grains in the colonized zone were partially or completely dissolved (Figure 3). Consistent with surface control, the partially dissolved mineral grains were dissolved along crystallographic and twinning planes. This is why the pathway stone is sugared. A subsample was embedded in epoxy, polished, and studied under SEM and electron dispersive spectroscopy (EDS). Secondary calcite and clay minerals were detected in pore spaces next to the colonized zone (Figure 4). Unlike the primary dolomite grains, which is composed equal amounts of Ca and Mg, the secondary calcite contained little Mg (Figure 4, EDS spectra). The clay minerals were identified on the basis of their composition, which includes Al and Si as well as Fe and Ca. (Figure 4, EDS spectra).

**Figure 2.**
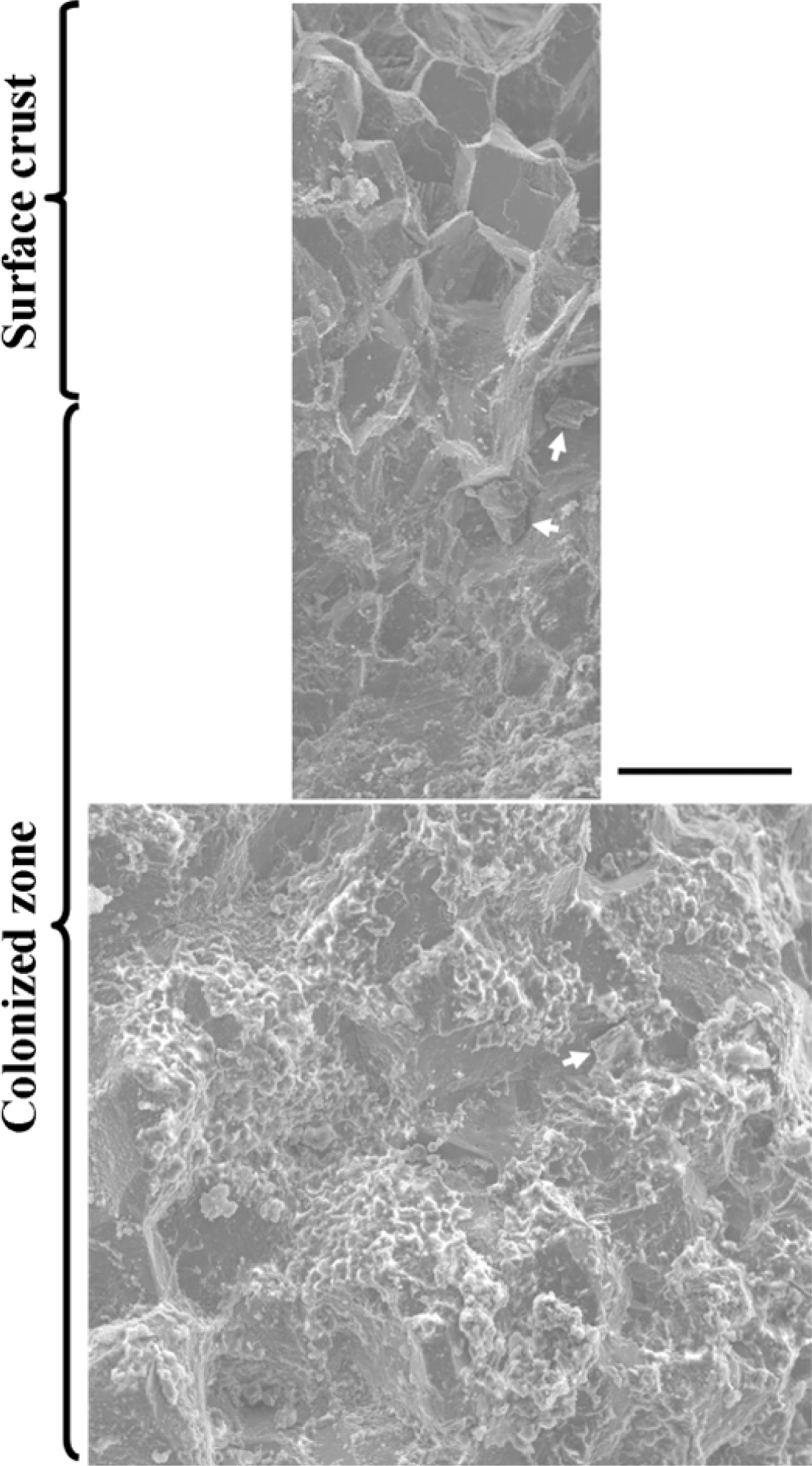
Scanning electron micrograph of an exfoliation crust, showing the smooth surfaces of mineral grains in the uncolonized outer layer and the biofilm in the colonized interior. Scale bar 200 µm.

**Figure 3.**
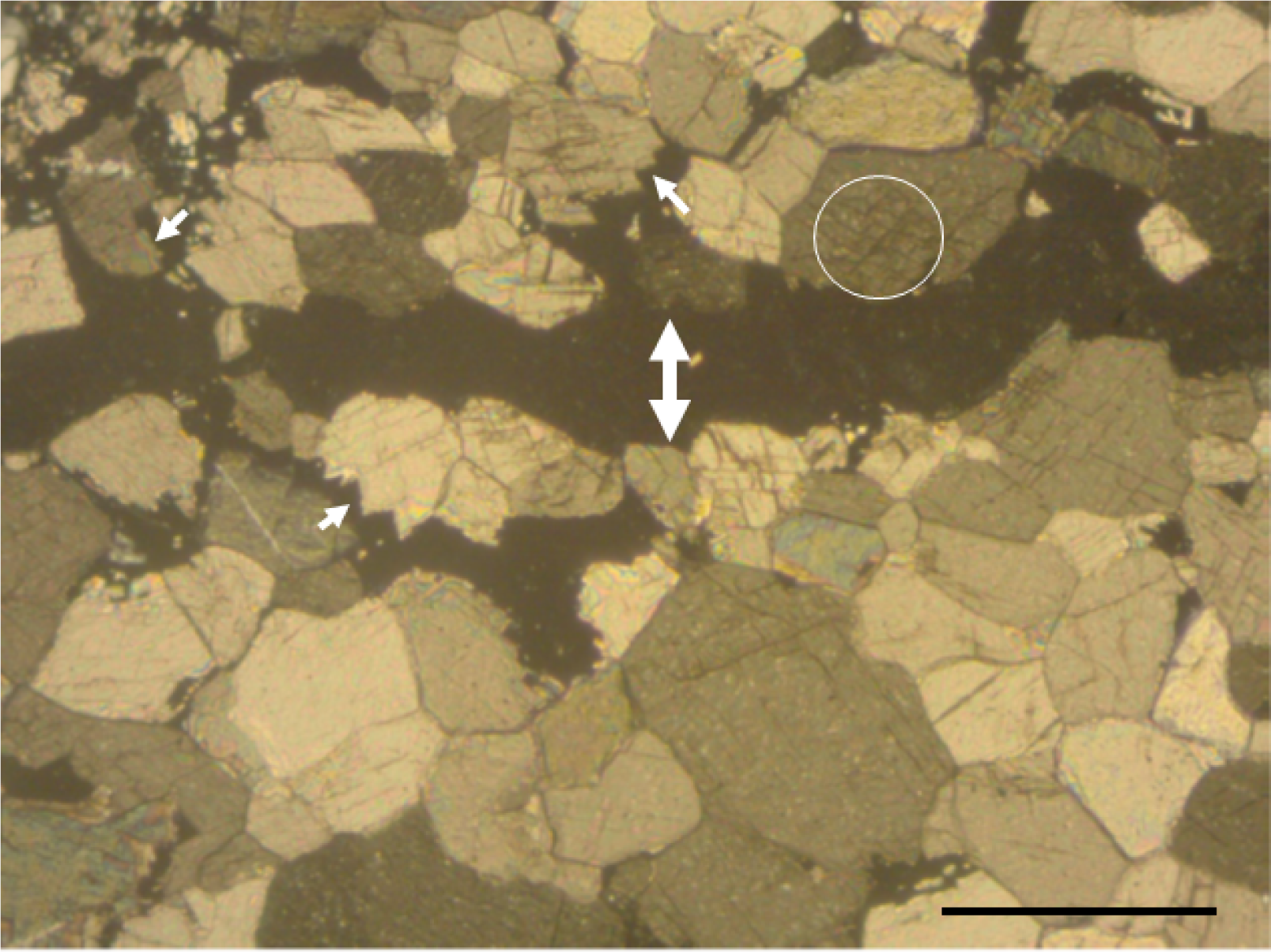
Polarized light micrograph of an exfoliation crust in petrographic thin section, showing the completely dissolved (two-sided arrow) and partially dissolved mineral grains in the colonized zone (arrows. Note that the partially dissolved mineral grains were dissolved along crystallographic (arrows) and twinning boundaries (circle). Scale bar 250 µm.

**Figure 4.**
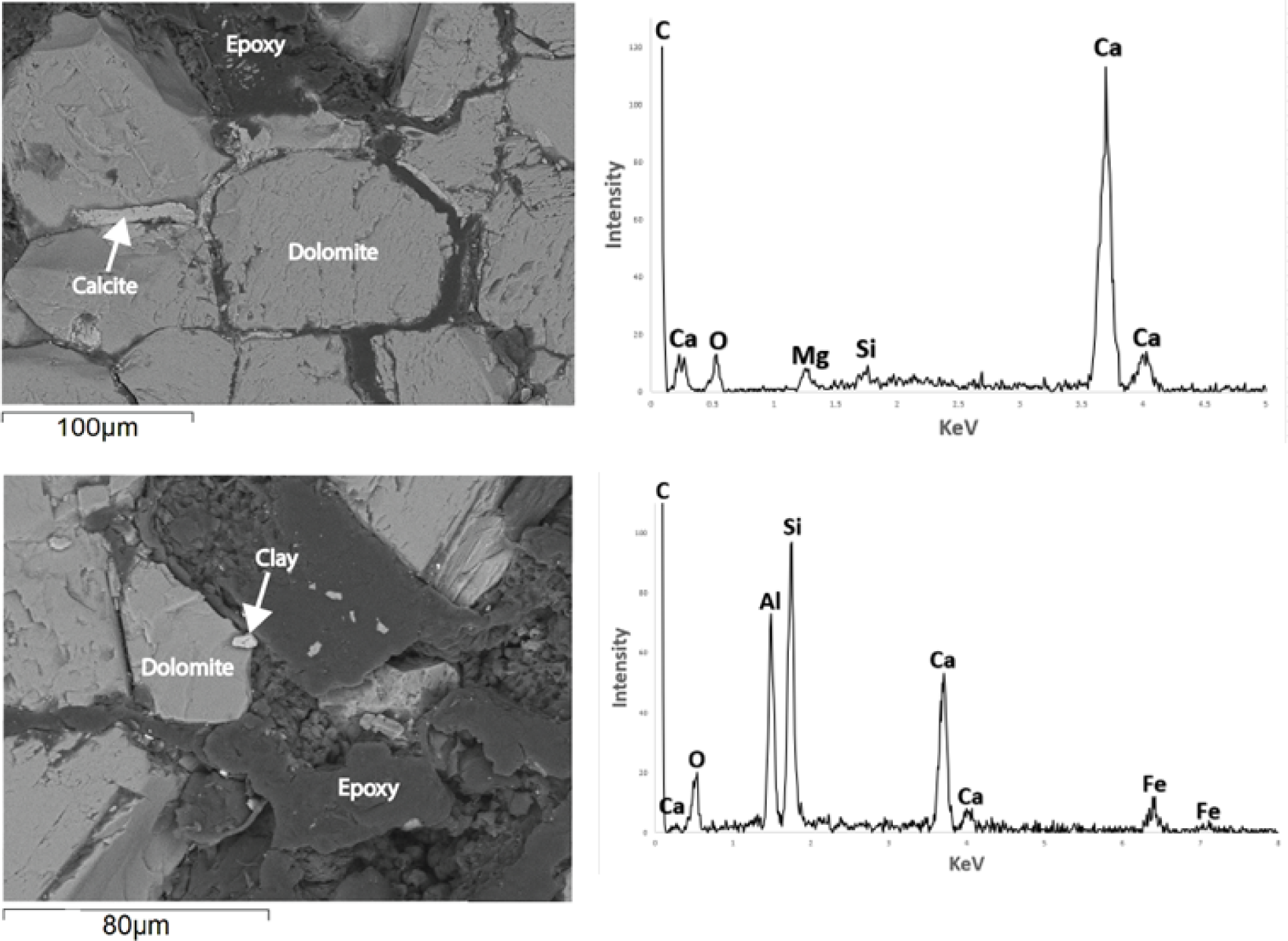
SEM micrograph of precipitates in the pore spaces in the outer uncolonized layer of an exfoliation crust and their EDS spectra, showing the presence of secondary calcite, composed largely of Ca, C, and O, with little Mg and Si (B), and clay minerals, composed of Si, Al, O, Fe, and Ca. These secondary minerals are evidence that calcium hydroxide is produced in the colonized zone below.

Four morphotypes of cyanobacteria were identified under the light microscope, including unicellular *Chroococcus* sp, *Aphanocapsa* sp, and *Stanieria* sp. and filamentous *Leptolyngbya* sp (Figure 5). DNA analyses detected eight cyanobacteria (Figure 6) and 12 heterotrophic bacteria. Of the eight cyanobacteria phylotypes, three (JQ404414, JQ4416, JQ404417) were related *Aphanocapsa muscicola*, *Stanieria cyanophaera*, and *Leptolyngbya frigida* by 92.3%, 94.5%, 93.8% in sequence similarity, respectively; together they account for 62.8 % of the 90 clones analyzed (Table 1). The other five, relatively rare cyanobacteria phylotypes (from JQ404418 to JQ404422) were related to *Leptolyngbya frigida*, *Stanieria cyanophaera*, *Aphanocapsa muscicola*, *Loriellopsis cavernicola*, and *Aphanocapsa muscicola* by 93.1%, 94.0%, 90.3%, 93.5%, and 90.9% in sequence similarity, respectively; together they account for 6.7% of our clone library (Table 1). Notably, the morphotype *Chroococcus* sp. was not present in our clone library. The 12 heterotrophic bacteria phylotypes (from JQ404423 to JQ404434), varying from 9.1% to 1.1% in abundance, were related, in varying sequence similarity from 85.6% to 99.8%, to *Mycoplasma alvi, Amaricoccous solimangrovi, Skermanella aerolata, Pontibacter humi, Oipengyuania sediminis, Orthithinibacillus contaiminans, Mesorhizobium carmichaelinearum, Sphingosinicella humi, Sphingomonas Altererythrobacter muriae, Sphingomonas oligophenolica*, and *Cultibacterium acnes* (Table 1).

**Figure 5.**
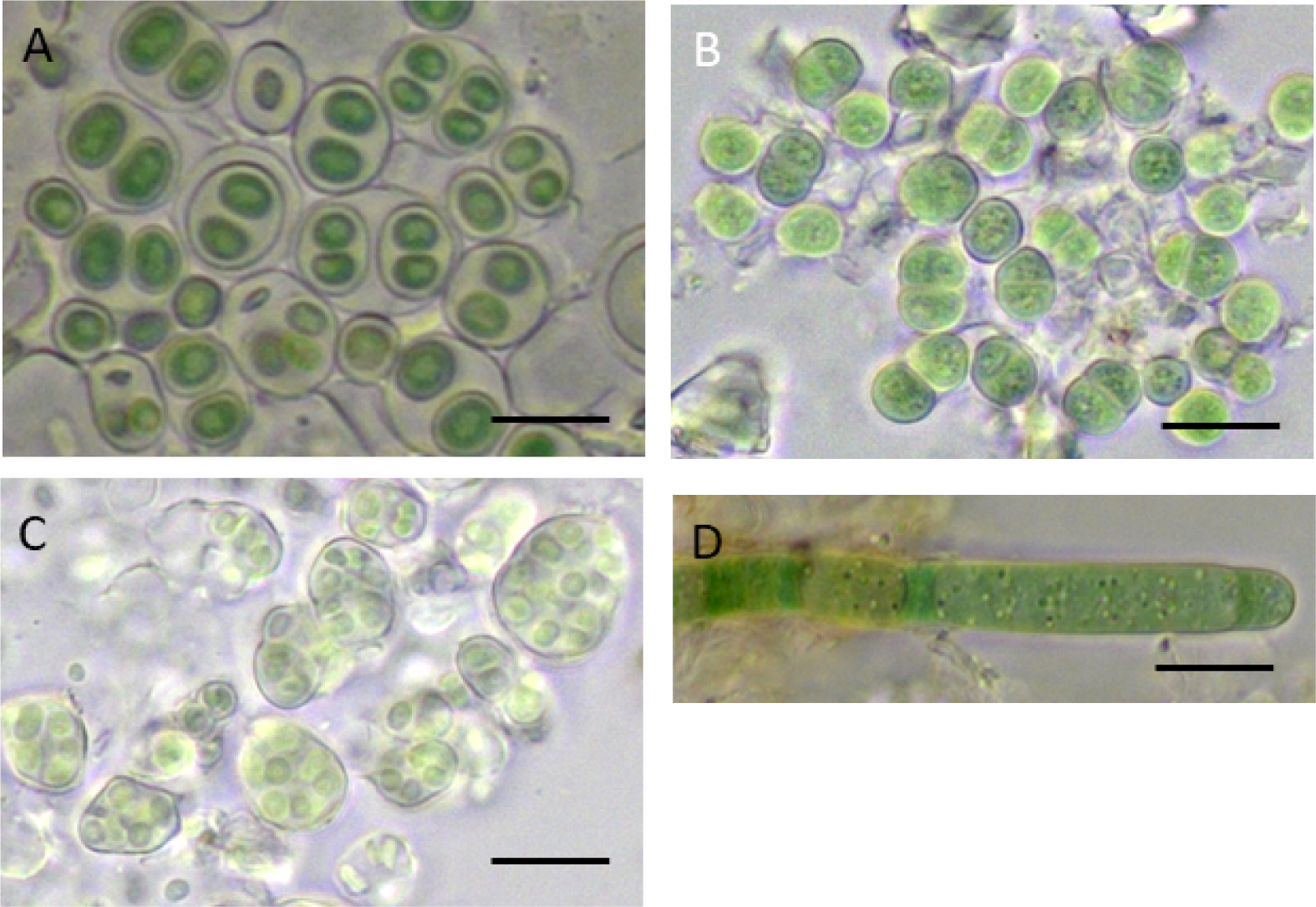
Light micrographs of endolithic cyanobacteria in Beijing. A) *Chroococcus* sp., B) *Stanieria* sp., C) *Aphanocapsa* sp, D) *Leptolyngbya* sp. Scale bar 5 µm.

**Figure 6.**
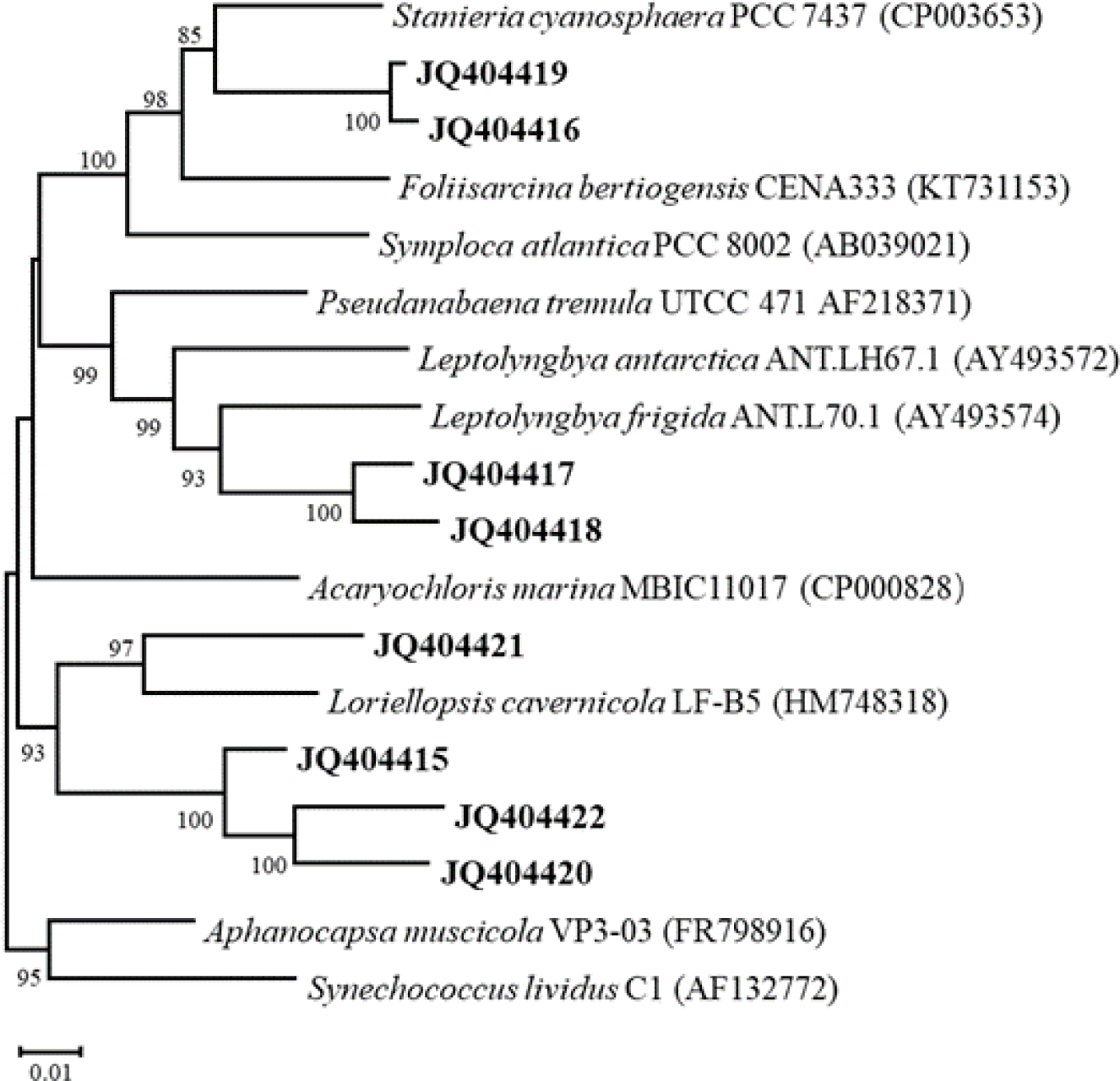
Phylogenetic tree of endolithic cyanobacteria in Beijing and close relatives.

**Table 1.**
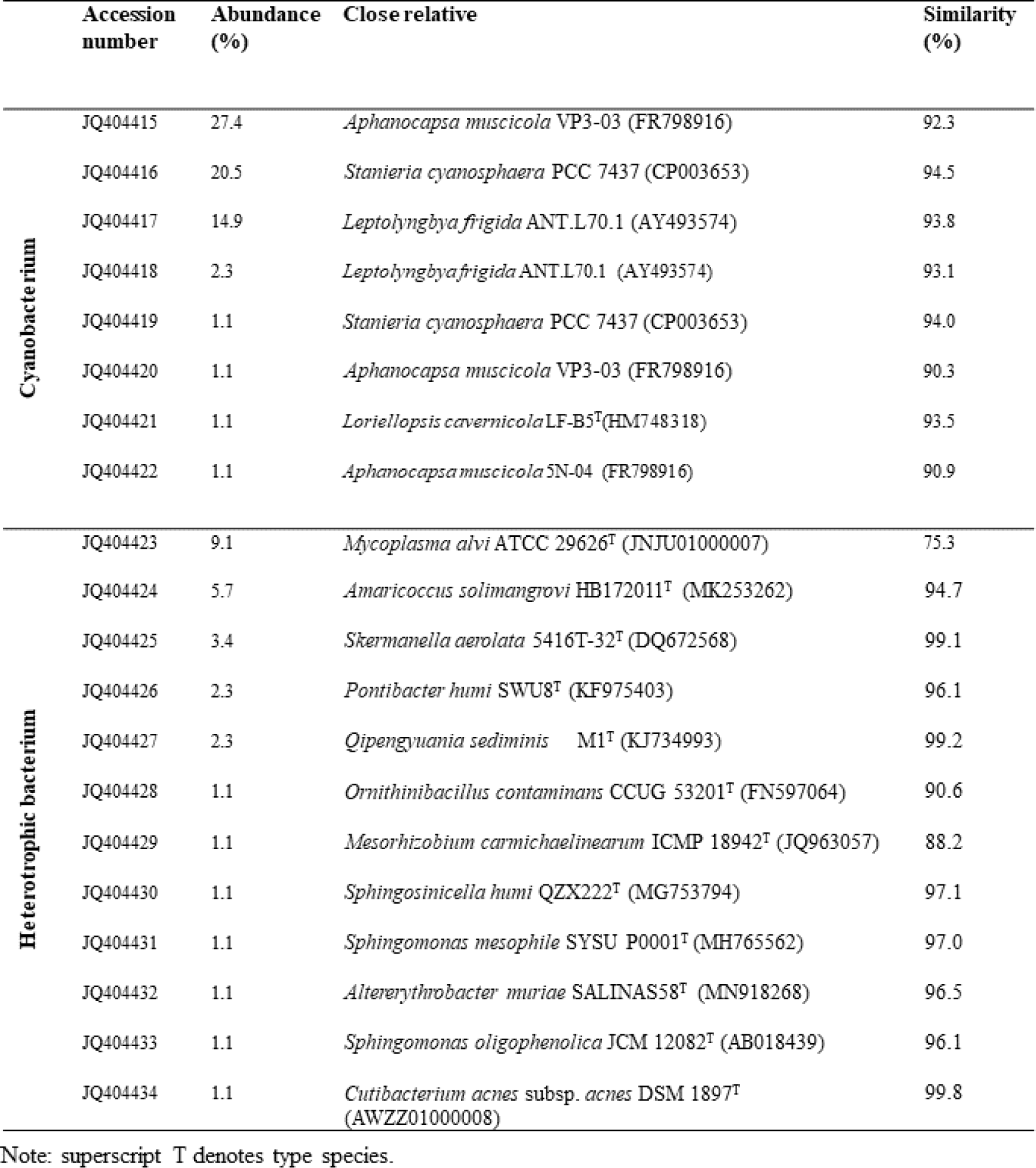
Cyanobacterial and bacterial 16S rRNA sequences recovered from colonized marble, their abundance (out of 90 clones), close relative, and sequence similarity to close relative.

## Discussion

Rainwater is slightly acidic, with pH around 6.5, due to dissolved CO_2_. The same water turns basic if the dissolved CO_2_ and resulting bicarbonate are rapidly removed. This chemistry was verified by Büdel and his colleagues in a cyanobacterial culture sandwiched in sand [27]. During the 2005/6 field season in Antarctica, one of us measured the midday interstitial water with pH papers. It was 8.5. Consequently, both the carbonate cement in the Mojave Desert sandstone and the silicate cement in the Antarctic sandstone are dissolved, resulting in stone decohesion and exfoliative weathering. Silicate-cemented sandstone transmits UV, whereas carbonate-cemented sandstone does not. Thus, the Antarctic sandstone also illustrates why iron oxides, which are virtually insoluble in aerobic circumneutral conditions, are destabilized and leached out. Colonized sandstone contains oxalic acid [54]. In the presence of organic acids and UV, iron oxides are reductively dissolved [55, 56]. While most of the iron is leached to the stone interior, a small amount is precipitated on the stone surface, blocking UV. A crust takes years or longer to form, however, so colonized stone is inevitably exposed after a crust exfoliates. Consequently, the top of the iron accumulation zone in the fractured stone is sharp and parallel to the stone surface (Figure 1B, upper inset). While this iron chemistry is most prominent in silicate-cemented sandstone, there are no reasons why it cannot take place on the UV irradiated surface of other rock types.

The stones in Beijing display all known hallmarks of biological weathering. Weathered marbles are associated with cracks and fissures (Figure 1C, D, E) and harbor endolithic cyanobacteria (Figure 1C, Figure 2). Colonized mineral grains are dissolved (Figures 3). Encrustation is accompanied by iron dissolution and reprecipitation, by formation of secondary calcium carbonates, and by formation of clay minerals (Figure 4). Stones that are horizontal and may be covered in snow lack a crust and instead disintegrate grain by grain, causing the partially dissolved mineral grains to be exposed, imparting a sugared appearance. Due to the exudate’s ability to dissolve silicates, such stones are self-cleansing of adhered dust as well (Figure 1E).

Endolithic cyanobacteria also exist in limestones in supralittoral and intertidal zone, where some species were recognized as euendolithic, capable of active carbonate dissolution or microboring [57, 58]. Recently, Garcia-Pichel and colleagues [59] developed an ingenious laboratory system to investigate microboring in *Mastigocoleus testaruma*, a filamentous euendolithic cyanobacterium that bores into solid calcite [59]. Microboring occurs under light, not in the dark, suggesting that it is an active process. Microboring is halted in the presence of Ca-ATPase inhibitors, a result that Garcia-Pichel et al. interpreted as indicating that microboring is coupled to active calcium excavation. According to them, calcite was dissolved by proton secreted from the leading end of the boring filament, while the dissolved calcium was taken up by the same cell and transported out of the borehole through the filament. In our opinion, calcite was dissolved by water, while the required Ca-ATPase activity has a more likely, alternative explanation. Ca ion is chaotropic, i.e. toxic in high concentrations [60]. The Ca-ATPases are needed to expel excess calcium leaked from the surrounding interstitial water. Inhibiting this activity leads to calcium poisoning. Indeed, as Garcia-Pichel et al. [59] themselves observed, the dissolved calcium precipitated upon exiting the borehole, suggesting it was highly alkaline and incompatible with cellular physiology. It could only have exited the borehole in the interstitial space and passively down a concentration gradient, not through the living organism.

Although euendolithic cyanobacteria are able to create their own pore spaces in the laboratory, they recruit noneuendolithic cyanobacteria when grown in their natural habitat [33, 34, 57, 61, 62]. The community in Beijing is no exception (Figures 5, 6, Table 1). Of the eight cyanobacteria phylotypes detected, only one is related to a known euendolith, *Leptolyngbya* sp. [63]. The growth of endolithic microorganisms in stone is limited by the availability of pore spaces. The euendolithic and noneuendolithic members may be in a mutualistic relationship where the former trade pore spaces for two critical services from the latter. One service is UV shielding during growth in cracks and fissures and after the surface crust exfoliates. In those vulnerable places or times, sheathed noneuendolithic cyanobacteria fortify their sheath with scytonemin, a brownish UV-screening pigment [53, 64-66]. Indeed, exposed cyanobacteria are brown or black (Figure 1D), shielding the sheathless, scytonemin-negative euendolithic cyanobacteria like *Leptolyngbya*.

The noneuendolithic cyanobacteria may also help their euendolithic comrades produce a CO_2_ deplete environment essential to microboring. The noneuendolithic cyanobacteria are physically necessarily between the atmosphere and the euendolithic species. In other words, they are positioned to intercept CO_2_ from the atmosphere. The euendolithic cyanobacteria have no option but to dissolve carbonates and, as a result, produce more pore spaces. The noneuendolithic cyanobacteria are presumably also equipped with Ca-ATPases and allow the dissolved calcium to exit around them without harm.

An endolithic community that is optimized to degrade its rock substrate may sound counterintuitive, but it is essential for long-term survival. A desert is not a permanent habitat. On geological timescales, deserts disappear while new deserts appear elsewhere on the planet, sometimes on the other side of an ocean. Continued existence requires that new deserts be settled as soon as they appear, or else the end of a desert would spell extinction. In other words, long term survival requires a constant production of propagules by colonized stones. Indeed, desert cyanobacteria were detected in randomly sampled urban aerosols [43]. Tomb stones throughout North America are colonized, including in Boston, Massachusetts, a city several thousand kilometers from the closest desert (Sun, unpublished data).

In what is a well-documented but poorly understood feature, endolithic communities recruit diverse heterotrophic bacterial decomposers but keep their cell numbers low [67, 68] [69] [32, 70]. This is also the case in Beijing (Figures 5, 6, Table 1). This feature is related to the need to keep rock dissolution going at fastest rates possible. Dead cyanobacteria no longer contribute to carbonate hydrolysis and need to be evacuated to make room for new growth. Cyanobacteria contain peptidoglycans in their cell wall, a complex structural polymer, as well as nucleic acids, proteins, and polysaccharides. Their depolymerization requires many bacteria species working in concert [71]. Yet, a larger bacterial presence beyond this function would be counterproductive to stimulating carbonate hydrolysis. Heterotrophic bacteria do not absorb CO_2_. They produce CO_2_.

While we are the first to draw a mechanistic link between stone weathering and rock-inhabiting microorganisms, we are not the first to discover their presence in stone monuments. In fact, colonized stone monuments have been previously reported from around the world, including the Parthenon of Acropolis in Athens, Greece [72], the exterior walls of Old Cathedral Building of Tongeren, Belgium [73], the Pyramid of Caius Cestius in Rome, Italy [33], the exterior of buildings in Brazil, Columbia, and Peru [74], the Acropolis at the Ek’s Balam, Yucatan, Mexico [69], and the Bayon Temple at Angkor Thom, Cambodia [75], to name just a few. The widespread presence of endolithic microorganisms in non-desert climates is consistent with our hypothesis in that it goes to show that: 1) propagules of endolithic microorganisms are indeed ubiquitous in the environment, as some of us previously suggested [44], and 2) arid climates are not essential to existence; only unobstructed access to sunlight is.

The widespread presence of rock-degrading endolithic microorganisms in urban landscapes challenges the generally accepted notion that rapid deteriorating stones in such settings are a consequence of exposure to acid rain [3, 7, 76-78]. Lichens are extreme sensitive to air pollution [79, 80], so deteriorating stones in cities may have more do with the fact that they are devoid of lichens and thus available to endolithic colonists and less or even nothing to do with acid rain. Beijing is a case in point. The city is situated in a region that is frequently exposed to aeolian dust, the region being proximal and downward to deserts in the northern and northwestern China [81]. The resulting carbonate aerosol, by the virtue of their high surface area, effective neutralize SO_2_ and NOx, the acidifying gases in air pollution [82]. But Beijing remains uninhabitable to epilithic lichens. The absence of a significant acid rain contribution to stone deterioration is evident in stones that are lichen-free but not yet colonized by endolithic cyanobacteria. These surfaces are sound and show no signs of deterioration (Figure 1E).

The deteriorating stone monuments in England and in the continental United States may not be unlike the situation in Beijing, but for a different reason. Although metropolises in those countries were plagued by notorious smoggy days in the 1940s and 1950s, by the 1980s and 1990s, when the sorry condition of stone monuments was noticed, air quality, especially SO_2_, had vastly improved [83]. This is not to say that no acid rain occurred, but it may not have been severe enough to be responsible for the observed stone deterioration. The fact that limestone deterioration at St Paul’s cathedral in London was occurring at the virtually same high rates from 2000 to 2010 as it was from 1980 to 1990 is suggestive [84]. We have begun to revisit some of the deteriorating stone monuments in the continental United States to see if they have been colonized by endolithic microorganisms. Stones visited so far, which include a cemetery near Los Angeles, the Field Museum in Chicago, and the Harvard Bixi in Boston, are all indeed colonized (Sun, unpublished data).

## Acknowledgements

This work was supported by NASA (grant# NNX08AO45G).

